# EEG power spectra and subcortical pathology in chronic disorders of consciousness

**DOI:** 10.1101/695288

**Authors:** Evan S. Lutkenhoff, Anna Nigri, Davide Rossi Sebastiano, Davide Sattin, Elisa Visani, Cristina Rosazza, Ludovico D’Incerti, Maria Grazia Bruzzone, Silvana Franceschetti, Matilde Leonardi, Stefania Ferraro, Martin M. Monti

## Abstract

**Objective:** To determine (i) the association between long-term impairment of consciousness after severe brain injury, spontaneous brain oscillations, and underlying subcortical damage, and (ii) whether such data can be used to aid patient diagnosis, a process known to be susceptible to high error rates.

**Methods:** Cross-sectional observational sample of 116 patients with a disorder of consciousness secondary to brain injury, collected prospectively at a tertiary center between 2011 and 2013. Multimodal analyses relating clinical measures of impairment, electroencephalographic measures of spontaneous brain activity, and magnetic resonance imaging data of subcortical atrophy were conducted in 2018.

**Results:** In the final analyzed sample of 61 patients, systematic associations were found between electroencephalographic power spectra and subcortical damage. Specifically, the ratio of beta-to-delta relative power was negatively associated with greater atrophy in regions of the bilateral thalamus and globus pallidus (both left > right) previously shown to be preferentially atrophied in chronic disorders of consciousness. Power spectrum total density was also negatively associated with widespread atrophy in regions of the left globus pallidus, right caudate, and in brainstem. Furthermore, we showed that the combination of demographics, encephalographic, and imaging data in an analytic framework can be employed to aid behavioral diagnosis.

**Conclusions:** These results ground, for the first time, electroencephalographic presentation detected with routine clinical techniques in the underlying brain pathology of disorders of consciousness and demonstrate how multimodal combination of clinical, electroencephalographic, and imaging data can be employed in potentially mitigating the high rates of misdiagnosis typical of this patient cohort.

**Search terms:** disorders of consciousness, subcortical pathology, EEG, MRI.

## Introduction

When patients survive a severe brain injury, but fail to fully recover the two cardinal elements of consciousness (i.e., awareness and arousal; (Laureys, 2005; Monti, 2012)), they are said to enter a disorder of consciousness (DOC; (Monti, Laureys, & Owen, 2010)). This label describes a set of conditions including coma, a state of unarousable unresponsiveness, the vegetative state (VS), a state of arousal in the absence of (self)awareness, and the minimally conscious state (MCS), a state in which patients can demonstrate some level of (self)awareness, albeit inconsistently. MCS can be further subdivided into MCS+, to indicate patients capable of high-level behavioral responses such as command following and intelligible verbalizations, and MCS-, to indicate patients only demonstrating low-level behaviors such as visual pursuit and appropriate contingent behavior (e.g., smiling or crying in response to emotional stimuli; (Bruno, Vanhaudenhuyse, Thibaut, Moonen, & Laureys, 2011)).

Over the past 20 years, electroencephalography (EEG) and magnetic resonance imaging (MRI) have been increasingly employed to monitor neurological status (Brenner, 2005), residual cognitive function (Chennu et al., 2013; Monti et al., 2015; Schnakers et al., 2008), state of awareness (Crone et al., 2015; Demertzi et al., 2019; Owen et al., 2006), and potential for recovery (Bagnato et al., 2010; Crone, Bio, Vespa, Lutkenhoff, & Monti, 2018; Schnakers et al., 2019) in DOC patients. In the context of bedside EEG, analysis of the magnitude of oscillations at different frequencies (i.e., power spectrum analysis) has been shown to be capable of differentiating DOC patients from patients with severe neurocognitive disorder but no consciousness impairment (Leon-Carrion, Martin-Rodriguez, Damas-Lopez, Barroso y Martin, & Dominguez-Morales, 2008), as well as clinical categories of chronic DOC (i.e., VS, MCS), with depth of impairment correlating with slower, larger amplitude oscillations (Lechinger et al., 2013; Lehembre et al., 2012). While this technique is sensitive to different injury etiologies (Fingelkurts, Fingelkurts, Bagnato, Boccagni, & Galardi, 2013), it is blind to the underlying anatomical damage. At a theoretical level, the mesocircuit theory of recovery of consciousness after brain injury predicts a relationship between the slowing and amplitude of electrocortical oscillations and the degree of pathologic changes taking place after trauma, hypoxia, or multifocal ischemia (Schiff, 2016). While indirect evidence exists in support of this model, with *in vivo* and *post-mortem* work demonstrating a relationship between damage to thalamus, loss of thalamo-cortical structural connectivity, and depth of impairment (Adams, Graham, & Jennett, 2000; Lutkenhoff et al., 2015; Zheng, Reggente, Lutkenhoff, Owen, & Monti, 2017), there are no data directly connecting the patterns of EEG power spectra and subcortical damage in long-term DOC patients, a gap which is not only problematic for the clinician’s interpretation of the observed EEG data, but also hampers our ability to monitor, through an inexpensive, bedside, repeatable technique interventions and their effects. In what follows, we address, in a large cohort of patients with chronic DOC, the heretofore untested relationship between observed electrocortical rhythms, patterns of subcortical brain atrophy (including thalamus, brainstem, and basal ganglia), and clinical measures of awareness and arousal.

## Methods

### Participants

A convenience sample of 116 patients was recruited from a larger database (n=153) (Nigri et al., 2017; Rosazza et al., 2016; Rossi Sebastiano et al., 2015; Rossi Sebastiano et al., 2018) of chronic DOC patients with severe brain injury, who underwent a 1-week assessment at the Coma Research Centre (CRC) of the Neurological Institute C. Besta in Milan, Italy, between 2011 and 2013. The assessment included (i) clinical evaluation, including the Coma Recovery Scale Revised (CRS-R; (Giacino, Kalmar, & Whyte, 2004)), (ii) neurophysiological evaluation, including resting EEG, and (iii) neuroradiological assessment, including MRI (see Table 1).

**Table 1:** Analyzed sample summary statistics of demographic, clinical, and electroencephalographic data per diagnostic group. (Abbreviations: VS, Vegetative State; MCS-Minimally conscious State “minus”; MCS+, Minimally Conscious State “plus”; MPI, months post-injury; T, traumatic; NT, non-traumatic; H, hemorrhagic injury, I, ischemic injury, A, anoxic; Orom/Verb, oromotor/verbal; Communic, communication.) (See Table S1 for a summary of demographic and clinical variables for the excluded datasets and comparison with the analyzed sample.)

|  | VS | MCS- | MCS+ |
| --- | --- | --- | --- |
| Age (mean, SD, y) | 53 (±14.21) | 54 (±18.77) | 46 (±16.77) |
| MPI (mean, SD, mo) | 23 (±16.05) | 44 (±37.17) | 53 (±64.97) |
| Sex | 14F, 23M | 11F, 6M | 0F, 7M |
| Etiology | 12T, 9H, 1I, 1HI, 14TA | 5T, 7H, 1I, 0HI, 4TA | 3T, 3H, 1I, 0HI, 0TA |
| Coma Recovery Scale Revised (CRS-R) |  |  |  |
| Total score | 6.24 (1.01) | 9.24 (1.25) | 10.43 (1.90) |
| Auditory | 1.03 (0.50) | 1.41 (0.51) | 2.00 (1.15) |
| Visual | 0.76 (0.43) | 2.59 (0.71) | 2.43 (1.40) |
| Motor | 1.92 (0.28) | 2.12 (0.33) | 2.14 (0.90) |
| Orom/Verb | 1.00 (0.33) | 1.24 (0.56) | 1.43 (0.98) |
| Communic | 0.00 (0.00) | 0.12 (0.33) | 0.57 (0.53) |
| Arousal | 1.54 (0.56) | 1.76 (0.56) | 1.86 (0.90) |
| Electroencephalography (Total power & frequency relative power) |  |  |  |
| Total power (μV<sup>2</sup>) | 138.22 (140.35) | 229.20 (261.81) | 178.96 (148.15) |
| Delta (1–4Hz) | 49.46% (13.46%) | 41.51% (14.35%) | 42.57% (16.97%) |
| Theta (4–8Hz) | 30.21% (11.09%) | 34.09% (9.73%) | 33.23% (11.58%) |
| Alpha (8–13Hz) | 9.71% (5.64%) | 11.89% (6.08%) | 8.33% (4.48%) |
| Beta (13–30Hz) | 7.78% (7.09%) | 9.59% (8.22%) | 13.89% (17.23%) |
| Gamma (>30Hz) | 2.84% (3.00%) | 2.91% (3.21%) | 1.98% (1.63%) |

The acquisition of both resting EEG and structural MRI datasets constituted the inclusion criteria for the present study. Experienced raters independently assessed each patient 4 times with the Italian version of the CRS-R (Sacco et al., 2011). The best-recorded performance was used to classify patients as VS or MCS. As described below, 55 patients were discarded due to the low quality of the MRI data, following previously established procedures (Lutkenhoff et al., 2015). Specifically, 24 datasets were excluded because of excessive movement artifacts, 16 datasets were excluded due to software failure in segmenting subcortical structures, 13 datasets were excluded due to poor quality in the estimation of normalized brain tissue volume, and 2 subjects were excluded due to large regions of signal dropout artifacts (e.g., implants) preventing tissue segmentation (see Figure S1). Importantly, as discussed below, exclusions do not bias the analyzed sample as compared to the full cohort (see also Table S1). No patients were excluded on the basis of the EEG data. The final sample included 61 patients with DOC (median age=53 years, range age = 20-82 years, 36 males), 37 VS and 24 MCS. Of these, 33% (n = 20) had a traumatic brain injury (TBI) and 67% (n = 41) had non-traumatic etiology (non-TBI). In particular, among non-TBI patients, 30% (n = 18) suffered from anoxic brain injury and 37% (n = 23) from haemorrhagic and/or ischemic brain injury. The median disease duration at the time of the study was 24 months (range = 5-198 months). (See Tables 1 and S1). The local Ethics Committee approved all aspects of this research and written informed consent was obtained from the legally authorized representative of the patients prior to their inclusion in the study.

### Data acquisition and analysis

#### EEG data acquisition and processing

Patients underwent polygraphic recordings between 2 pm and 9 am on the following day (Rossi Sebastiano et al., 2018). Recordings included electrooculography (EOG), electromyography (EMG) from the sub-mental muscle, a bipolar precordial electrocardiogram (ECG) derivation and an impedance thoracic pneumogram. EEG recordings were made with Ag/AgCl surface electrodes (impedances *<*5kΩ) and acquired at a sampling rate of 256 Hz using a computerized system (Micromed SpA, Mogliano Veneto, Treviso, Italy). Raw EEG signals were recorded, against a common reference electrode to allow off-line data reformatting, using a 19 EEG electrode array placed according to the 10-20 International System (including frontal (Fp1, Fp2, F3, F4), central (C3, C4), parietal (P3, P4), and occipital (O1, O2) electrodes). We preliminarily divided the awake from sleep based on the polygraphic signal. Spectral EEG analysis was then performed on one epoch of 5-minute, consecutive, artifact-free, awake EEG signal. To minimize issues deriving from different levels of arousal, we selected the EEG epochs as close as possible to the beginning of the recording, in order to prevent signal degradation due to the prolonged recording and to maintain, as much as possible, the same reproducible characteristics between recordings in different patients. The selected epochs were filtered (1-70 Hz, 12 db/octave), followed by a 50 Hz notch filter to suppress the noise of the electrical power line, reformatted against the linked ear-lobe reference. In order to remove blink-artifacts, we applied an ICA-artifact rejection algorithm. Then, the selected EEG activity was divided into 90 non-overlapping 2 s segments and analyzed using the fast Fourier transform. Absolute total power and relative power were evaluated in the delta (1-4 Hz), theta (4-8 Hz), alpha (8-13 Hz), beta (13-30 Hz) and gamma (>30 Hz) bands, and averaged within each EEG channel.

#### MRI data acquisition and processing

Neuroimaging data were obtained with a 3T MRI machine (Achieva, Philips Healthcare BV, Best, NL; 32-channel coil). The MRI protocol included a high-resolution 3D-TFE T1-weighted sequence (185 sagittal slices, TR=9.781 ms, TE=4.6 ms, voxel size = 1 mm^3^, flip angle = 8°). Analysis of subcortical structures was conducted using a technique known as shape or vertex analysis, part of the FMRIB software library (FSL; FMRIB, Oxford, UK), following a previously established pipeline (Lutkenhoff et al., 2015; Schnakers et al., 2019). Briefly, MR images were brain-extracted using optiBET (Lutkenhoff et al., 2014), then the thalamus, caudate, putamen, globus pallidus, hippocampus, and brainstem were segmented using FSL FIRST (Patenaude, Smith, Kennedy, & Jenkinson, 2011), for each patient and structure separately, and reconstructed into 3-dimensional vertex meshes, as depicted in Figure 1. In addition, the normalized brain volume (NBV), a measure of global brain atrophy including white and gray matter volume, was calculated for each patient using FSL SIENA (Smith et al., 2002).

**Figure 1.**
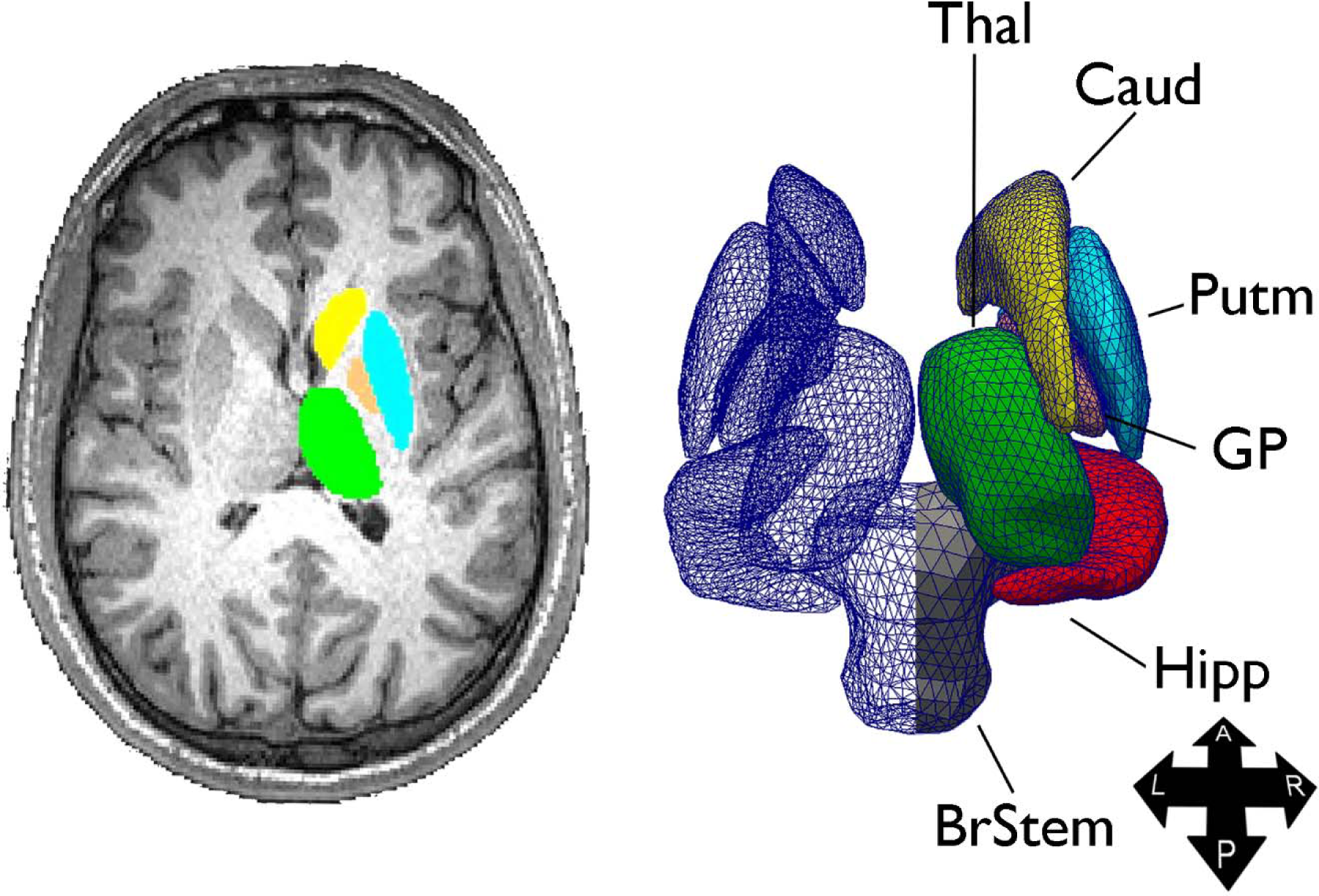
Sample structure extraction (left) and 3-dimensional triangle vertex mesh (right). (Abbreviations: A, anterior; BrStem, brain stem; Caud, caudate; GP, globus pallidus; Hipp, hippocampus; L, left; P, posterior; Putm, putamen; R, right; Thal, thalamus. Figure from (Lutkenhoff et al., 2015).)

#### Statistical Analyses

In what follows, we describe four analysis which were employed to assess the relationship between clinical/behavioral measures of impairment, electrophysiological recordings, and subcortical measures of atrophy.

#### EEG analysis

To assess the relationship between EEG spectral power and clinical grouping, we ran a mixed-model linear analysis with EEG (relative) power as the dependent variable, laterality (left, right, middle), channel position (Fp, F, C, P, O), and EEG features (total power, delta, theta, alpha, beta, gamma bands) as the repeated measures, diagnosis (VS, MCS-, MCS+), etiology (TBI, non-TBI), sex, EEG frequency, age, and months post-injury as fixed variables and subjects as the random variable. As described below, the significant interaction between EEG features and diagnosis was followed up with one mixed-model analysis per each EEG feature (using the same model, albeit without the EEG feature as repeated variable). Individual mixed-models were followed up with pairwise post-hoc comparisons between diagnostic groups (i.e., VS, MCS-, MCS+), with Šidák correction for multiplicity.

#### EEG – MRI analysis

In the second analysis, we related EEG spectral features to subcortical shape measures. However, because of significant correlations among spectral characteristics across electrodes and frequency bands, spectral data first were entered into a principal component analysis (PCA), with varimax rotation. The PCA returned 8 components with eigenvalue greater than 1, cumulatively explaining 91.8% of the total variance. The first three components presented a similar pattern, each loading negatively, in all electrodes, on the delta band and positively on beta, alpha, and theta bands, respectively (henceforth, *β*/*δ* ratio component, *α*/*δ* ratio component, and *θ*/*δ* ratio component, respectively). The fourth component loaded positively on all gamma frequency electrodes (henceforth, γ component), while the fifth component loaded positively on the total power for each electrode (henceforth, total power component). Finally, the last three components appeared to capture diffuse statistical covariance between electrodes, although with a preference for loading, respectively, positively on the delta band and negatively on the alpha band in right hemispheric electrodes (henceforth, *δ*/*α*_*RH*_ ratio component), loading positively on delta band and negatively on the alpha band in Fp electrodes (henceforth, *δ*/*α*_*Fp*_ ratio component), and loading negatively on both the alpha and theta bands in the parietal and occipital electrodes (henceforth, *αθ*_(P,O)_^−^). (See Figure S2 for a depiction of the strength of each component for VS, MCS-, and MCS+.) EEG components were then entered, as independent variables, in a general linear model predicting local shape patterns (e.g., atrophy). Sex, age, time-post-injury, etiology (i.e., TBI vs non-TBI), were included as covariates, along with NBV (to ensure that observed tissue displacement reflect local subcortical shape changes independent of overall brain atrophy (Lutkenhoff et al., 2015; Schnakers et al., 2019)). Group-level significance was assessed with a non-parametric permutation test at a level of *p <* 0.05 corrected for multiplicity using family-wise cluster correction and threshold free cluster enhancement (TFCE) as implemented in FSL randomize (Smith & Nichols, 2009).

#### CRS-R – MRI analysis

In this analysis, we related the patients’ behavioral presentation, as captured by the CRS-R subscales, with subcortical atrophy. Because of significant correlations between the subscales of the CRS-R (i.e., the desired independent variables), behavioral data were entered into a PCA performed analogously to the one described above. The analysis returned 3 components with an eigenvalue greater than 1, cumulatively explaining 69.57% of the variance. The three components loaded on, respectively, the auditory, visual, and arousal subscales (henceforth, audio-visual-arousal component), the motor subscale (henceforth, motor component), and the oromotor and communication subscales (henceforth, oromotor-communication component). As in the previous analysis, the three components were entered as independent variables in a general linear model predicting subcortical shape, along with the same covariates described above. Group-level significance was assessed identically to the previous analysis.

#### Predicting DOC level from EEG spectral features

Finally, we employed a binary logistic regression to evaluate the degree to which global atrophy and EEG measures related to diagnosis (i.e., VS vs. MCS). With a 3-block model, we attempt to predict diagnosis from a model including demographic variables only (i.e., age, sex, months post injury, etiology (TBI vs. non-TBI)), a model including demographics and a measure of global atrophy (including global white matter and gray matter), and a model including demographic, global atrophy, and EEG variables (i.e., the 8 EEG components).

## Results

### EEG results

The mixed-model analysis revealed a significant interaction (*F* (10, 1309.055) = 16.599, *p <* 0.001) between diagnostic group (i.e., VS, MCS-, MCS+) and EEG features (i.e., total power, delta, theta, alpha, beta, gamma frequency bands; see Figure 2 and Table 1), along with a significant main effect of diagnosis (*F* (2, 4389.124) = 5.158, *p* = 0.006), EEG features (*F* (5, 1309.055) = 6.368, *p <* 0.001), and months post-injury (*F* (2, 4220.566) = 5.407, *p* = 0.020). Follow-up mixed-model analyses (one per EEG feature) revealed a significant effect of diagnosis on total power (*F* (2, 594.374) = 19.115, *p <* 0.001; with pairwise-comparisons revealing that VS patients show significantly less total power than both MCS groups), delta (*F* (2, 770.006) = 4.620, *p* = 0.010; with VS patients having the most relative delta power, significantly more than MCS-patients), theta (*F* (2, 779.789) = 12.268, *p <* 0.001; with VS patients showing significantly less relative theta power than both MCS groups), alpha (*F* (2, 748.766) = 13.231, *p <* 0.001; with MCS-patients showing significantly more relative alpha power than VS and MCS+ patients, and VS patients showing greater relative alpha power than MCS+ patients), and gamma (*F* (2, 746.625) = 5.416, *p* = 0.005; with MCS+ patients showing significantly less relative gamma power than MCS- patients). (For further description of the EEG data in this cohort see (Rossi Sebastiano et al., 2018).)

**Figure 2.**
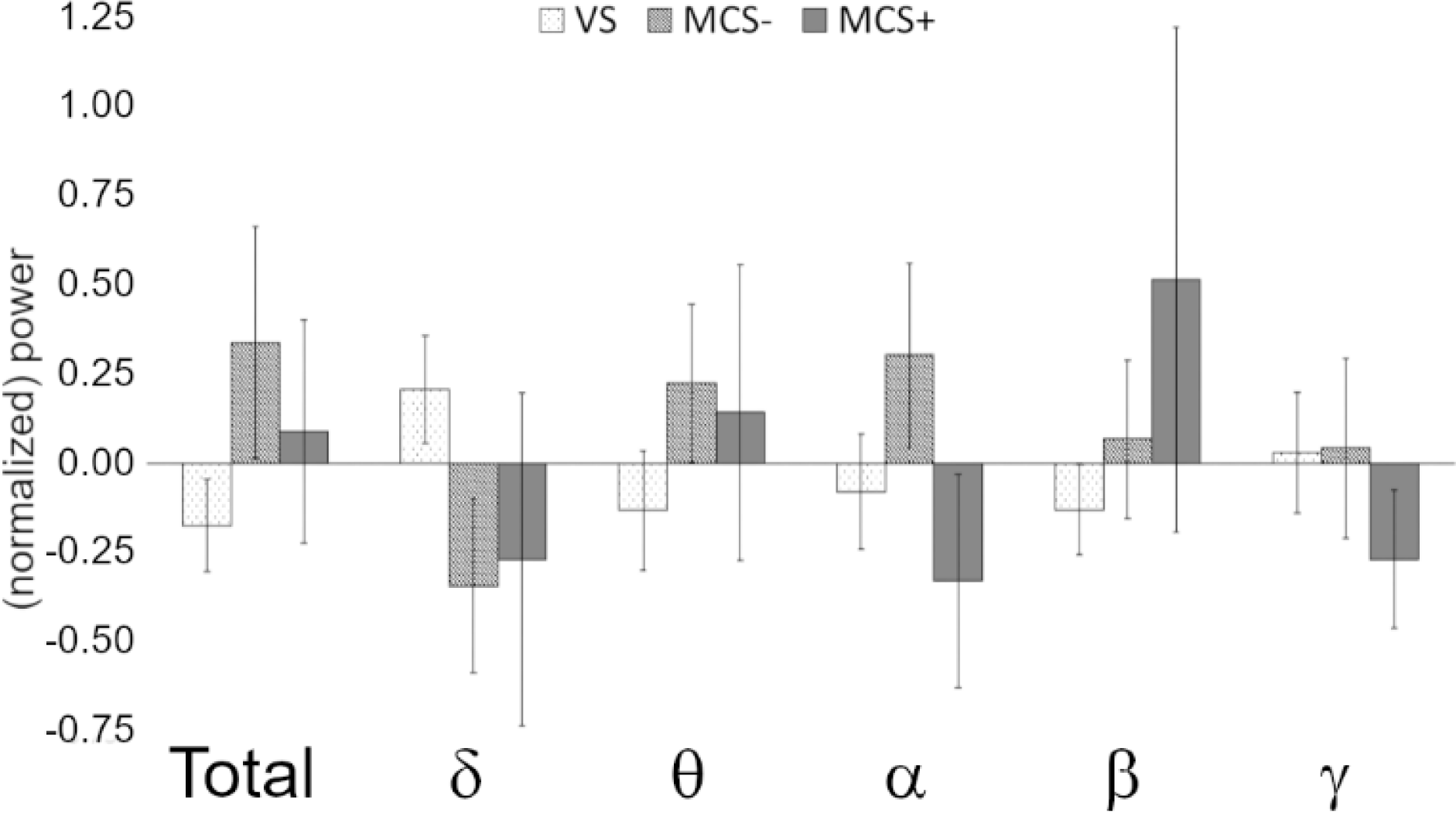
Summary of EEG data. Total and relative power at each frequency band (to allow displaying on the same axis, values are normalized within each variable). (Error bars represent standard errors.)

### EEG – MRI analysis

As shown in Figure 3, three of the EEG factors exhibited significant correlations with local atrophy measurements. Specifically, the *β*/*δ* ratio component was negatively associated with greater atrophy in bilateral thalamus (left: *t* = 3.28, *p* = 0.025, 1,007 significant vertices, (sig. vert.), right: *t* = 2.85, *p* = 0.041, 544 sig. vert.), bilateral globus pallidus (left: *t* = 4.26, *p* = 0.002, 491 sig. vert.; right: *t* = 3.42, *p* = 0.025, 356 sig. vert.), left caudate (*t* = 3.37, *p* = 0.02, 793 sig. vert.), and right hippocampus (*t* = 3.21, *p* = 0.02, 735 sig. vert.) (see Fig. 3a). The *θ*/*δ* component was negatively associated with increased atrophy in right putamen (*t* = 3.72, *p* = 0.037, 140 sig. vert.) and right globus pallidus (*t* = 3.89, *p* = 0.037, 30 sig. vert.) (see Fig. 3b). Finally, the total power component was negatively associated with widespread atrophy in the brainstem (including the 4^th^ ventricle region; *t* = 4.41, *p* = 0.004, 3704 sig. vert.), as well as the left globus pallidus (*t* = 4.00, *p* = 0.008, 365 sig. vert.) and right caudate (*t* = 5.48, *p* = 0.018, 193 sig. vert.) (see Fig. 3c). No significant associations were detected for the remaining components.

**Figure 3.**
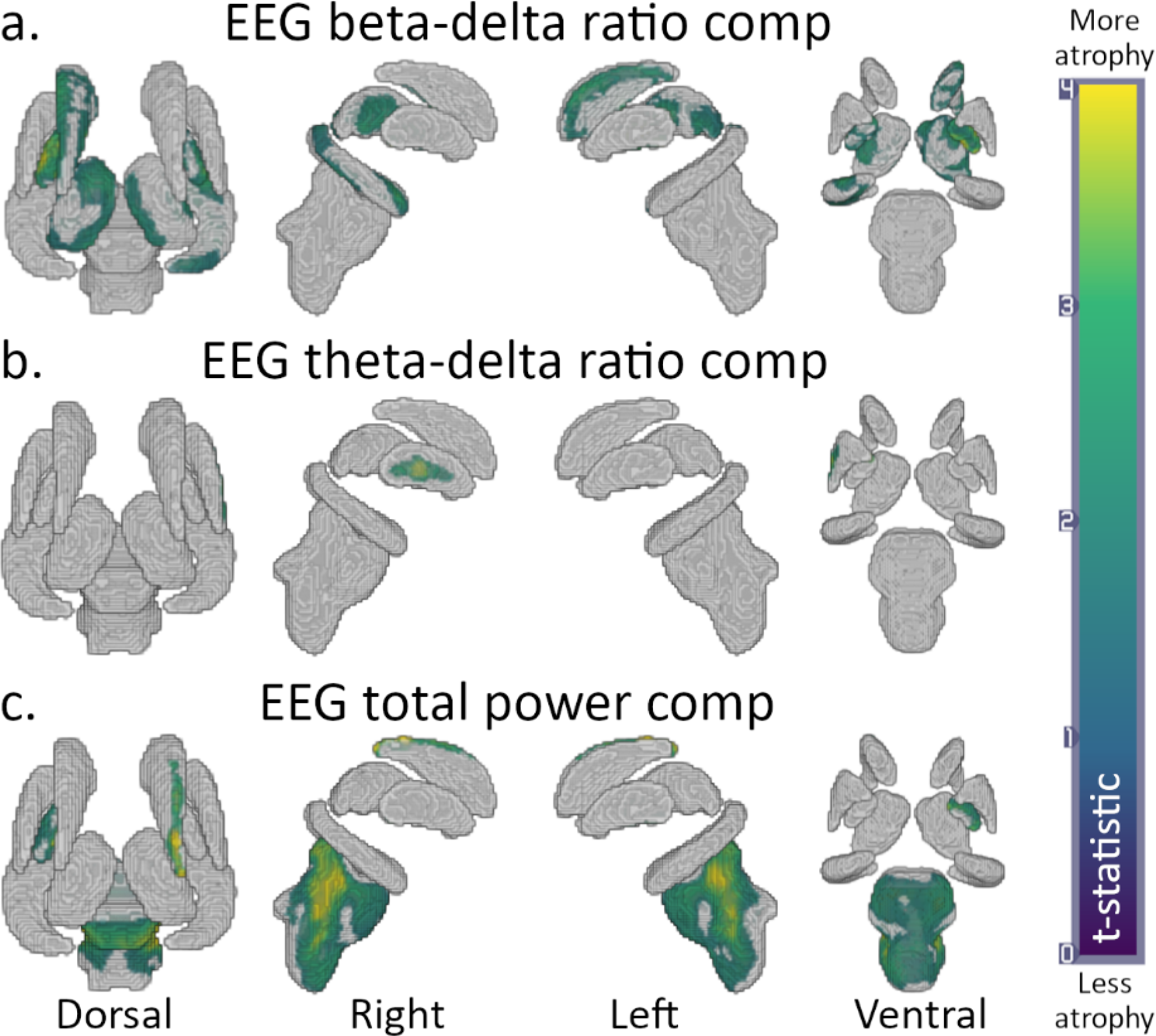
Results for the rest EEG – MRI analysis. (a) Results for the β/δ ratio component; (b) results for the θ/δ component; (c) results for the total power component. Warmer colors indicate regions with greater atrophy; gray indicates no significant effect.

### CRS-R – MRI analysis

As shown in Figure 4a,b, two of the CRS-R components exhibited significant correlations with local atrophy measurements. Specifically, the motor component was negatively associated with greater atrophy in broad regions of the brainstem (*t* = 3.69, *p* = 0.074, 4,301 sig. vert.; see Fig. 4a) while the oromotor-communication component was negatively associated with greater atrophy in regions of the brainstem (*t* = 2.74, *p* = 0.05, 943 sig. vert.), left putamen (*t* = 3.23, *p* = 0.023, 712 sig. vert.), and right globus pallidus (*t* = 2.38, *p* = 0.049, 108 sig. vert.).

**Figure 4.**
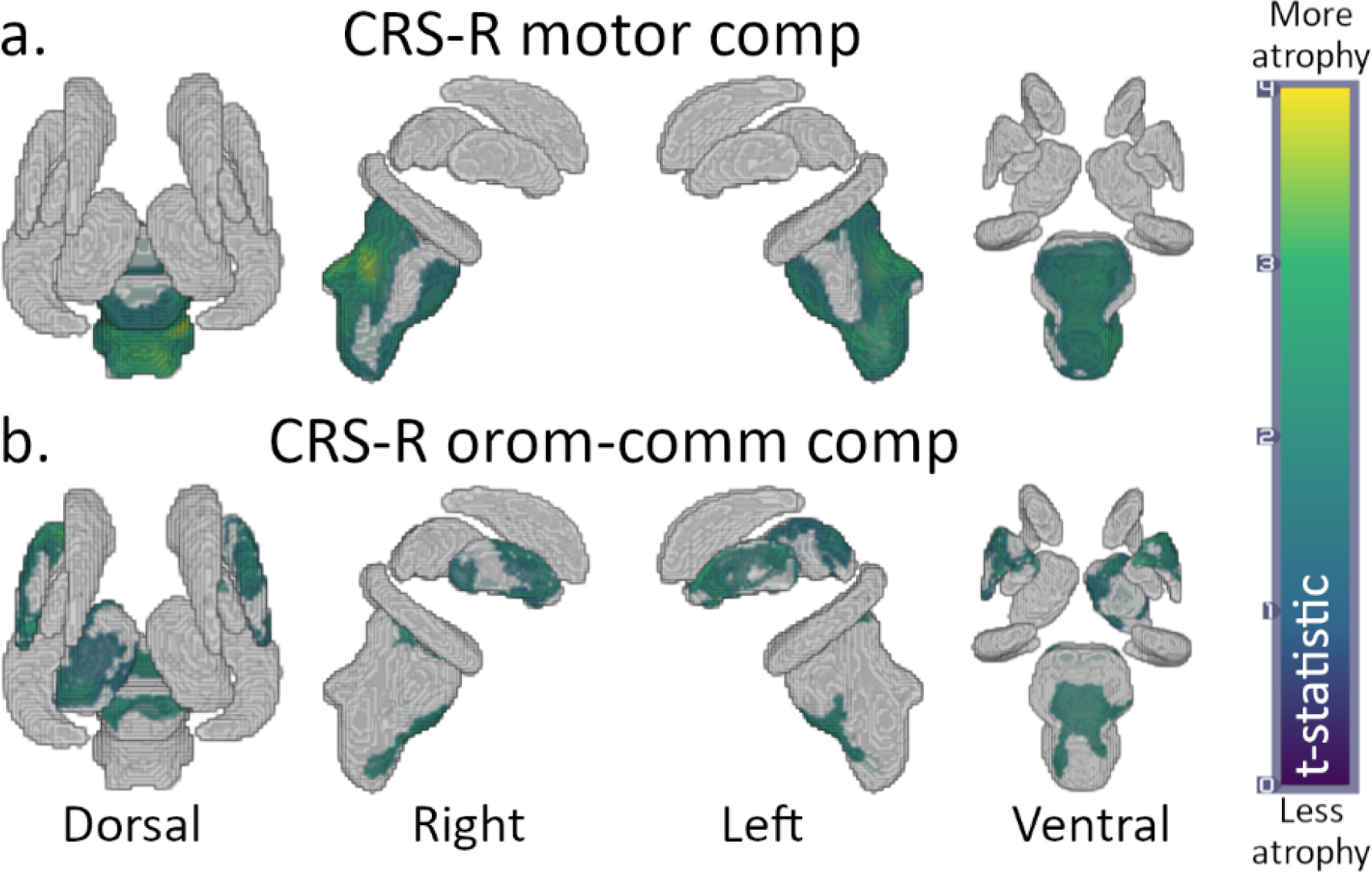
Results for the CRS-R – MRI analyses. (a) Results for the CRS-R motor component; (b) results for the CRS-R oromotor-communication component. (See Fig. 3 for color interpretation.)

### Predicting DOC level from EEG spectral features

As shown in Figure 5a and Table 2, diagnosis (i.e., VS vs. MCS) was predicted significantly better by the model including EEG components, overall brain atrophy (including both white matter and gray matter), and demographic components (i.e., age, sex, etiology (TBI vs. non-TBI), and months post injury; χ^*2*^(54) = 29.09, *p <* 0.001, Nagelkerke pseudo-*R*^*2*^_*Nag*_ = 0.51), compared to the model with demographics and brain atrophy (χ^*2*^(58)=16.66, *p <* 0.001, *R*^*2*^_*Nag*_ = 0.32), as well as the model with demographics only (χ^*2*^(59) = 8.93, *p* = 0.003, *R*^*2*^_*Nag*_ = 0.18). Indeed, the full model achieved better performance (i.e., area under the curve, *AUC* = 0.87), sensitivity and specificity (sens/spec; 0.79, 0.81, respectively) than both other models (*AUC* = 0.78, sens/spec 0.58/0.78 and *AUC* = 0.67, sens/spec 0.29/0.89 for the demographics and atrophy and the demographic only models, respectively). Notably, the contribution of increasingly complex models (i.e., adding brain atrophy and EEG components) is to increase the model’s sensitivity to MCS (at the cost of a loss of specificity).

**Table 2:** Binary logistic regression results. Top: comparison of the area under the curve, sensitivity, specificity, and precision of the model with demographic components only, the model with demographic components and the global brain atrophy factor, and the model with demographic components, global brain atrophy factor, and EEG components. Bottom: Individual predictors selected for the full model (i.e., DEM+Brain+EEG). (Abbreviations: area under the curve, AUC; sensitivity, sens.; specificity, spec.; precision, prec.)

| Model | df | $\Delta X^2$ | p | $R^2$ | AUC | Sens | Spec | Prec |
| --- | --- | --- | --- | --- | --- | --- | --- | --- |
| Intercept | 60 |  |  |  |  |  |  |  |
| DEM | 59 | 8.93 | 0.003 | 0.18 | 0.67 | 0.29 | 0.89 | 0.64 |
| DEM+Brain | 58 | 16.66 | <0.001 | 0.32 | 0.78 | 0.58 | 0.78 | 0.64 |
| DEM+Brain+EEG | 54 | 29.09 | <0.001 | 0.51 | 0.87 | 0.79 | 0.81 | 0.73 |
| 95% CI |  |  |  |  |  |  |  |  |
| Parameter | b | SE | $\beta$ | OR | z | p | LB | UB |
| (Intercept) | -1.92 | 0.50 | -0.79 | 0.15 | -3.86 | <0.001 | -2.89 | -0.95 |
| DEM: Sex-Age | 0.05 | 0.02 | 1.44 | 1.06 | 3.15 | 0.002 | 0.02 | 0.09 |
| Normalized Brain | 1.27 | 0.37 | 1.27 | 3.56 | 3.44 | <0.001 | 0.55 | 1.99 |
| EEG: Total power | 0.66 | 0.28 | 0.66 | 1.94 | 2.39 | 0.017 | 0.12 | 1.21 |
| EEG: $\delta/\alpha(RH)$ | 0.63 | 0.57 | 0.63 | 1.88 | 1.11 | 0.269 | -0.49 | 1.75 |
| EEG: $\theta/\delta$ | 0.61 | 0.32 | 0.61 | 1.84 | 1.88 | 0.060 | -0.03 | 1.25 |
| EEG:<br>$\alpha\theta(P,O)[Neg.]$ | 0.59 | 0.36 | 0.59 | 1.80 | 1.65 | 0.100 | -0.11 | 1.29 |

**Figure 5.**
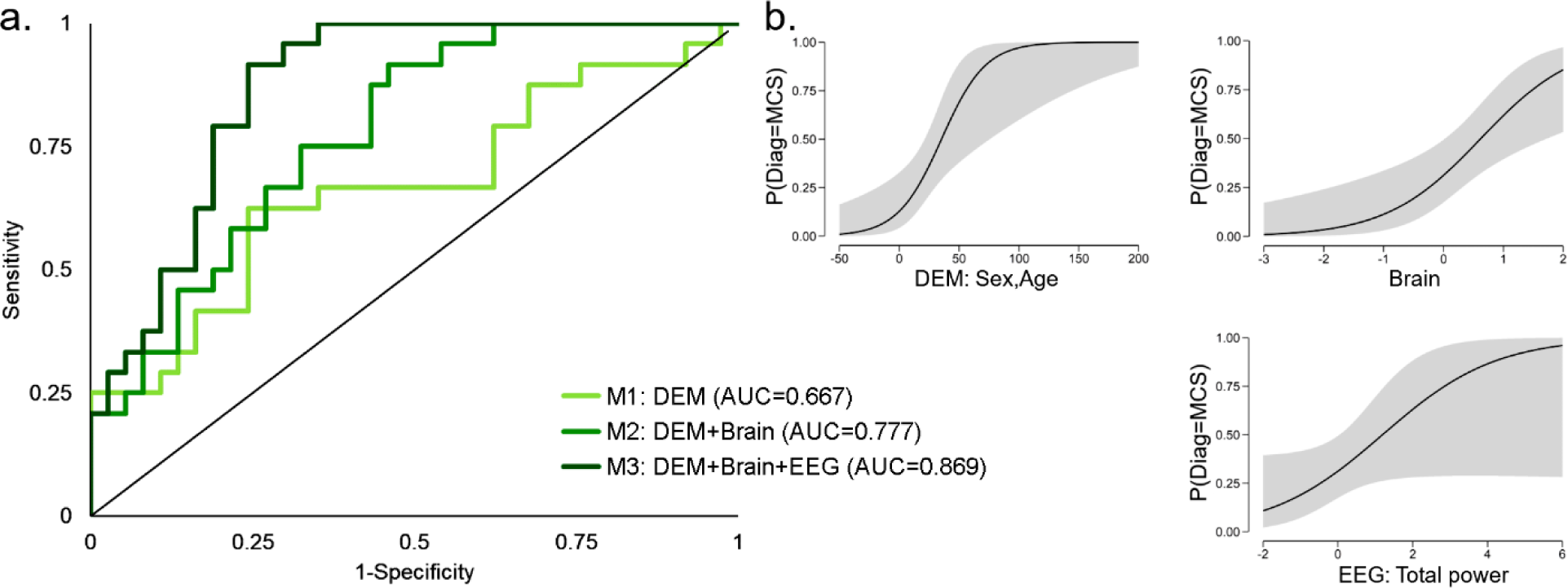
(a) ROC curve for the three binary logistic models classifying patient diagnosis (i.e., VS versus MCS) on the basis of demographic components alone (M1), demographic components and overall brain atrophy (M2), and demographic components, overall brain atrophy, and EEG components (M3). (b) Conditional estimate plots for each of the significant variables selected in the full model; top row: Demographic (age, sex) component, brain overall atrophy (i.e., NBV); bottom row: EEG total power component, EEG theta/delta component).

Finally, in terms of individual variables, as shown in Figure 5b and Table 2, the full model selected one demographic component (age/sex; *β* = 1.44, *OR* = 1.06, *p* = 0.002), overall brain atrophy (*β* = 1.27, *OR* = 3.56, *p <* 0.001), as well as the EEG total power (*β* = 0.66, *OR* = 1.94, *p* = 0.017).

## Discussion

We report three main findings addressing the relationship between severity of the impairment of consciousness, EEG spectral profile, and subcortical atrophy. First, our results show that spectral profiles recorded with conventional EEG map onto specific patterns of subcortical brain pathology (as observed with MRI), in line with theoretical proposals (Schiff, 2010; Schiff, 2016). Indeed, we find that the ratio of fast (i.e., beta) to slow (i.e., delta) frequencies component is negatively related to atrophy in thalamic regions well known to be associated with severity of impairment after brain injury, as shown in *post-mortem* (Adams et al., 2000) and *in vivo* (Lutkenhoff et al., 2015; Lutkenhoff et al., 2013) studies. Damage within thalamus, along with functional (Monti et al., 2015) and/or structural (Zheng et al., 2017) disconnection of thalamo-cortical projections, is indeed central to current theories of recovery from severe brain injury (Schiff, 2010), with greater de-afferentiation leading to a state of functional thalamic quiescence and, through cell-intrinsic mechanisms (Crunelli et al., 2018), delta oscillations (Schiff, 2016; Schiff, Nauvel, & Victor, 2014). Indeed, interventions aimed at up-regulating thalamic activity have been shown capable of enhancing behavioral responsiveness in DOC patients (Schnakers & Monti, 2017). The origin of beta frequency oscillations in the thalamo-cortical circuit remains unclear (Koelman & Lowery, 2019), nonetheless, there are at least two lines of evidence connecting them to DOC. On the one hand, a lesser state of thalamic de-afferentiation might result, through circuit-level interactions, in a thalamic bursting component expected in the beta frequencies (Schiff et al., 2014). On the other hand, beta oscillations have been associated with the globus pallidus (and subthalamic nucleus), a region which we find strongly negatively associated (particularly in its lateral segment), with the beta/delta component, and which has been previously implicated in electrocortical and behavioral arousal in animal models (Qiu, Vetrivelan, Fuller, & Lu, 2010) and behavioral arousal in several pathologies including DOC (Lutkenhoff et al., 2015). Some evidence suggests that beta oscillations can be induced by increased inhibition of the globus pallidus (Mirzaei et al., 2017), which, in DOC, would imply a lesser dysfunction within the mesocircuit (i.e., greater inhibitory input from the striatal medium spiny cells towards the pallidum; Schiff 2010). Thus, whether of thalamic or pallidal origin, or a superposition of the two, beta oscillations in DOC might mark a comparatively less disconnected mesocircuit.

Second, similarly to results obtained in a different (large) cohort of chronic DOC patients (Lutkenhoff et al., 2015), subscales of the CRS-R mapped onto different underlying patterns of atrophy, with oromotor/verbal and communication subscales component loading on left thalamic and bilateral putamen, as well as subregions of brainstem, and the motor component loading mainly on extensive brainstem atrophy. While the present results are similar to those reported previously (Lutkenhoff et al., 2015), there also are important differences. For example, no association was found between clinical measurement and brainstem atrophy in our previous work, and the pattern of thalamic atrophy in the two studies is not fully overlapping. It is difficult, however, to fully evaluate the meaning of these differences since the PCA components extracted in the two studies loaded differently on each CRS-R subscale. Therefore, although we do find a consistent set of regions correlating with CRS-R communication subscale in the two studies, spanning left thalamus and putamen, the details of the associations between all subscales and subcortical brain pathology remain to be fully characterized. Third, consistent with prior work (Corchs et al., 2019; Estraneo et al., 2016; Lechinger et al., 2013), we find that EEG spectral features are relevant to diagnosing a patient’s chronic state of consciousness (i.e., VS versus MCS), a decision known to be susceptible to a relatively high misdiagnosis rate (Monti & Owen, 2010; Monti et al., 2015; Schnakers, Giacino, Kalmar, Piret, & Lopez, 2006; Schnakers et al., 2009). Specifically, EEG components, together with demographic information, could correctly classify patients across the conscious/unconscious line (as behaviorally defined) with ∼87% success, leveraging on demographic information (i.e., sex, age), overall brain atrophy, and EEG features (i.e., total power and *θ*/*δ* components). It is also noteworthy that in our data combination of EEG and MRI data is additive in terms of enhancing discriminability across groups, strengthening the idea that multimodal approaches are a desirable way of assaying different – and complementary – aspects of DOC pathology (Coleman, Bekinschtein, Monti, Owen, & Pickard, 2009; Schiff, 2006) and, more broadly, brain function and health (Kumral et al., 2020; Nentwich et al., 2020), particularly in the context of deriving predictive models from brain data (Engemann et al., 2020). Despite the good concordance between behavior-based diagnosis and the classification based on demographic, brain atrophy, and EEG variables, the two approaches still disagree over one third of the cases. Seven MCS patients were classified as being in VS and seven VS patients were classified as being MCS. The MCS patients classified as VS by our analysis could reflect the fact that behavioral diagnoses compress very different EEG profiles (for example, patients with traumatic or anoxic etiology) into the same clinical category. Similarly, the VS patients classified as MCS by our algorithm could either be a reflection of the variance in the spectrum of oscillations that are compatible with a state of unconsciousness, or a genuine misdiagnosis (Edlow et al., 2017; Monti et al., 2015; Owen et al., 2006).

Finally, in evaluating the above results, the reader should be mindful of some limitations. First, our results are skewed by survivor bias effects; we might thus be representing a spectrum of impairment which, while severe, excludes the even greater damage present in patients who do not survive later than a year post injury. The same is true with respect to the fact that 55 datasets had to be excluded from the collected sample. While it would have been ideal to be able to retain more of the sample, conventional quality control limited our ability to analyze the full dataset. Nonetheless, the final analyzed sample is similar to that of other MR-based work in the field (e.g., (Demertzi et al., 2015; Monti, Vanhaudenhuyse, et al., 2010; Stender et al., 2014; Vanhaudenhuyse et al., 2010)). Furthermore, as shown in Table S1, the analyzed sample and the excluded sample are matched on almost all demographic and clinical variables, suggesting that our procedure, while affecting statistical power, is not introducing additional bias to the collected sample. Second, while VS and MCS-subgroups exhibit the expected imbalance across male and female patients (cf., (Lutkenhoff et al., 2015; Monti, Vanhaudenhuyse, et al., 2010; Stender et al., 2014)), it should be noted that the MCS+ subgroup is entirely composed of male patients, thus inviting caution in extrapolating results exclusive to this subgroup (e.g., differences in relative power distribution). Second, unexpectedly, we observe that MCS+ patients exhibit lower alpha relative power compared to MCS-patients. Nonetheless, considering that the two groups are matched in terms of overall relative power across alpha and beta frequencies (24.4% and 24.2% for MCS-and MCS+, respectively), it is possible that the decreased relative power in the alpha band observed in MCS+ is simply the converse of the increased relative power observed in the beta band (both relative to MCS-). Although caution should be exercised in making such inferences from a small subset of our data, it is conceivable that MCS-patients could have a more “rigid” rhythmic EEG less susceptible to variations as compared to MCS+ patients. Third, due to significant correlations across channels within and across power bands, in order to perform the regression analyses presented above, we had to first reduce the independent variables by means of a PCA. While conventional, it does affect the interpretation of our results in as much as we cannot directly assess whether the effects we report in mixed component (e.g., the *β*/*δ* component) are principally due to either frequency or to their combination. Nonetheless, each of the three ratio components correlates strongly with the “raw” ratio of each pair of frequencies (calculated by taking the ratio of the average relative power across all channels; specifically, r=0.97, r=0.87, and r=0.85 for *β*/*δ, α*/*δ*, and *θ*/*δ*, respectively) as well as with each numerator and denominator variable (albeit numerically more with the nominator for *β*/*δ* and *α*/*δ* components; see Table S2). This suggests that our results can be reasonably interpreted as reflecting actual ratios in relative power. Third, gamma frequencies are known to often contain residual muscle artifacts. We decided to keep them in the analysis mainly because, even if they do contain artifacts, including this component still contributes to explaining variance in the signal. Had we not included it, any variance across patients due to motion would have *de facto* been subsumed by the unexplained variance term. In this sense, our approach is analogous to the conventional inclusion of motion parameters in functional MRI analysis. Fourth, while we report associations between brain damage in subcortical regions and EEG spectral features, this does not necessarily imply that the pinpointed areas are themselves the generators of specific oscillatory rhythms at rest.

Finally, it should also be pointed out that the present work did not include an additional independent sample to confirm the classification results, thus inviting caution in the extrapolation of the results towards new cohorts. As multi-site efforts increase, larger cohorts of high-quality data will permit full application of such approaches.

In conclusion, this work bridges different levels of analysis of patients surviving severe brain injury, uniting DOC brain pathology (Adams et al., 2000; Lutkenhoff et al., 2015; Lutkenhoff et al., In press), clinical evaluation (Giacino et al., 2004), and power spectral features (Brenner, 2005; Fingelkurts et al., 2013; Schnakers et al., 2019) in a framework that makes use of routine multimodal clinical data and informs both basic science and the diagnostic process. Furthermore, the present data also show that the pattern of association between spectral profile, brain damage, and clinical variables observed in the acute setting (Schnakers et al., 2019) persist through the chronic timeframe, in line with the idea that brain injury is best thought of as a long-term disease as opposed to an “event” (Masel & DeWitt, 2010).

## Author Contributions

SF, MMM, ML, MGB conceived the study; DRS, SF, EV, oversaw collection and clinical interpretation of EEG data; LD, MGB, AN, CR oversaw collection and clinical interpretation of MRI data; ESL, AN, MMM, carried out the analysis presented in the paper; ESL, AN, SF, MMM, interpreted the results and drafted the manuscript; all authors provided critical feedback on the manuscript and revised the manuscript; SF, MMM, MGB secured funding.

## Ethical standards

The authors assert that all procedures contributing to this work comply with the ethical standards of the relevant national and institutional committees on human experimentation and with the Helsinki Declaration of 1975, as revised in 2008.

## Supplementary Online Materials

**Table S1.**
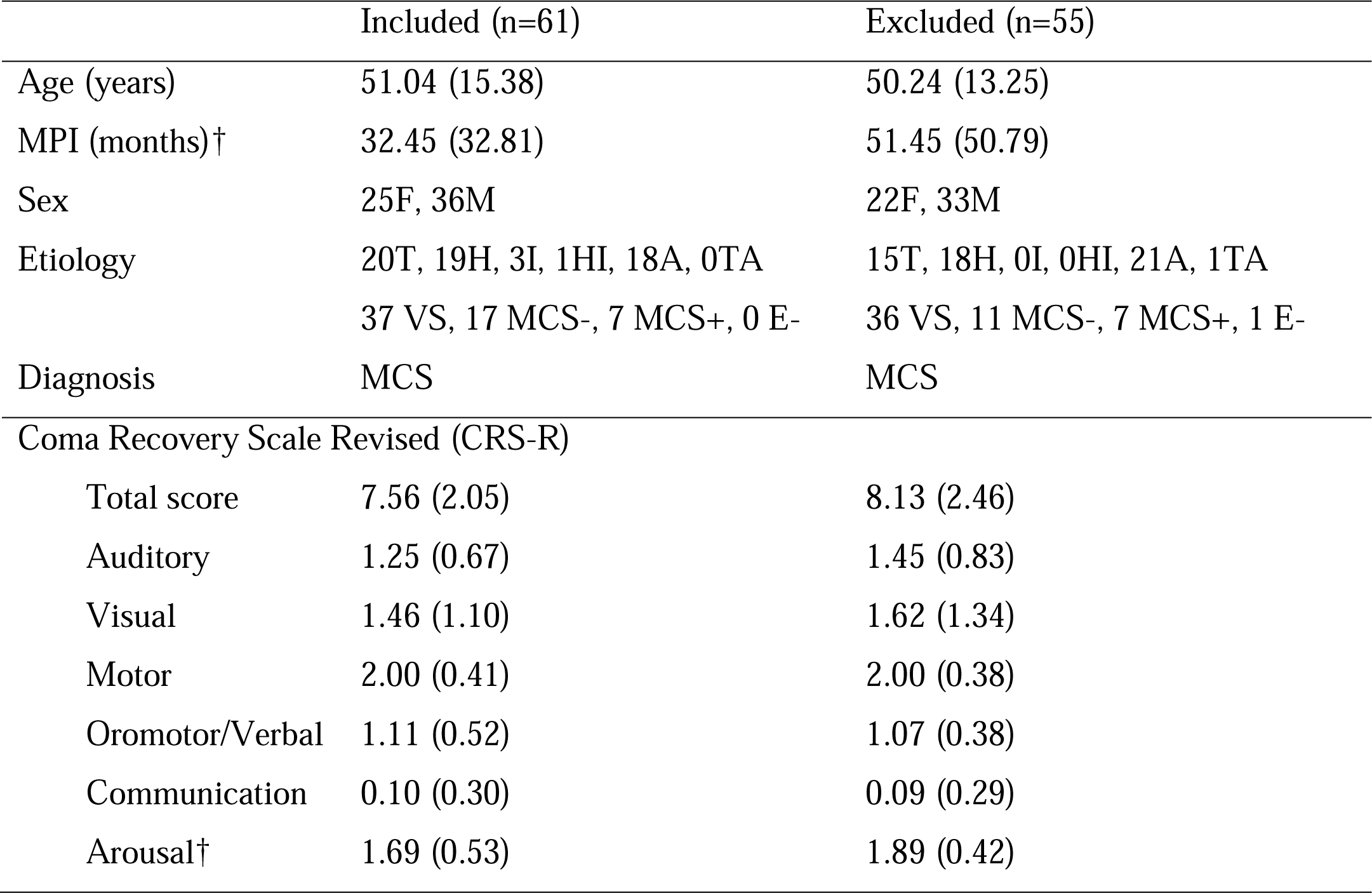
Comparison of demographic and clinical variables of patients included in the final analyzed sample (n-61) and patients excluded during the MRI processing/QC step (n=55). (Abbreviations: VS, Vegetative State; MCS-Minimally conscious State “minus”; MCS+, Minimally Conscious State “plus”; MPI, months post-injury; T, traumatic; H, hemorrhagic injury, I, ischemic injury, A, anoxic. ‘†’ indicates significant difference across groups prior to correction for multiple comparisons. No significant differences were observed after correction for multiplicity.)

**Table S2.** Spearman correlation between the three ratio components (β/δ, α/δ, and θ/δ) and the raw ratio of each pair of frequencies (calculated by taking the ratio of the average relative power across all channels), as well as the numerator frequency (β, α, and θ, respectively) and the denominator frequency (δ for all components).

|  | raw ratio | numerator | denominator |
| --- | --- | --- | --- |
| $\beta/\delta$ comp | 0.95 | 0.94 | -0.61 |
| $\alpha/\delta$ comp | 0.87 | 0.94 | -0.78 |
| $\theta/\delta$ comp | 0.85 | 0.79 | -0.80 |

**Figure S1.**
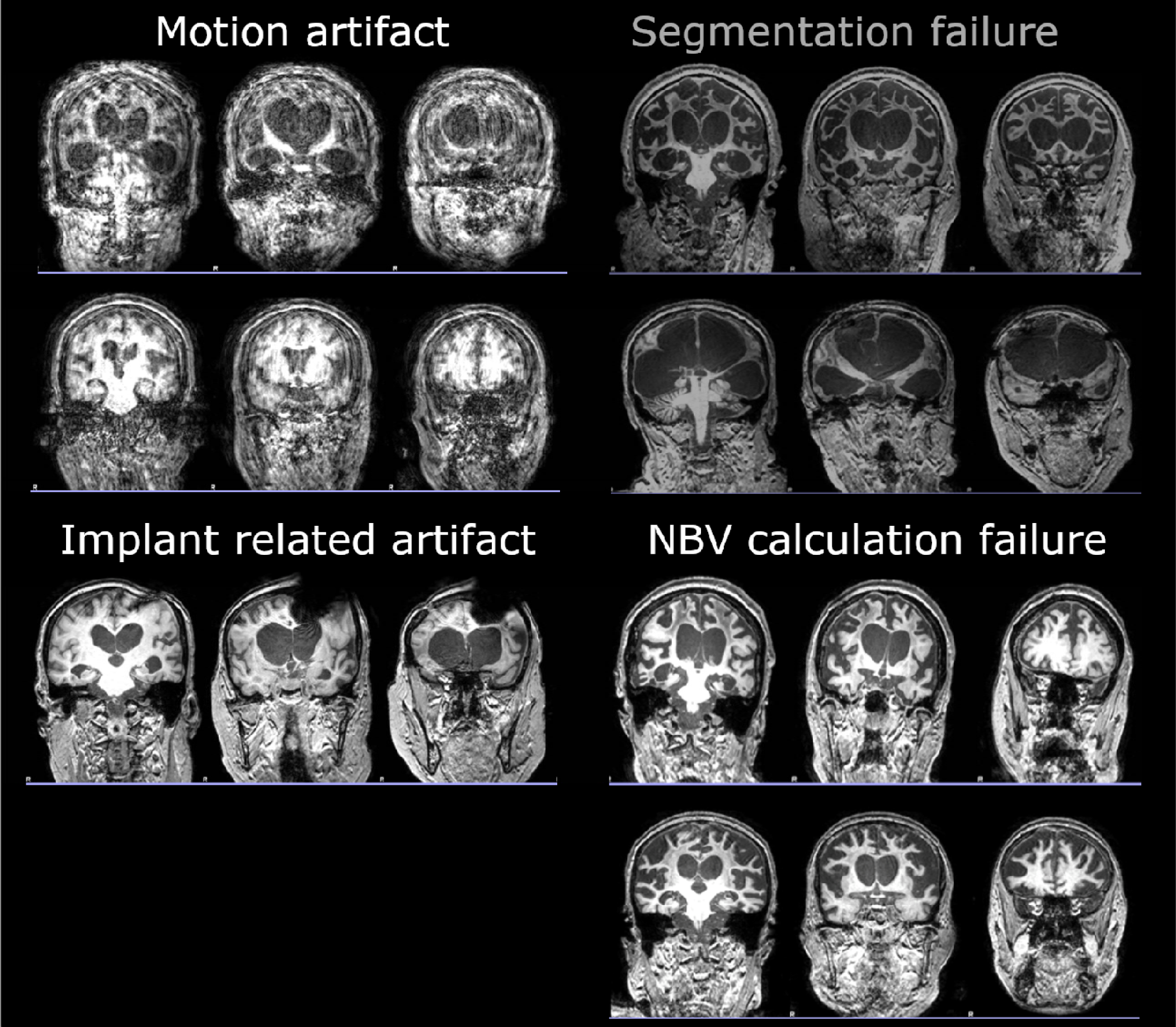
Examples of excluded dataset (for each category) during the MRI quality control process.

**Figure S2.**
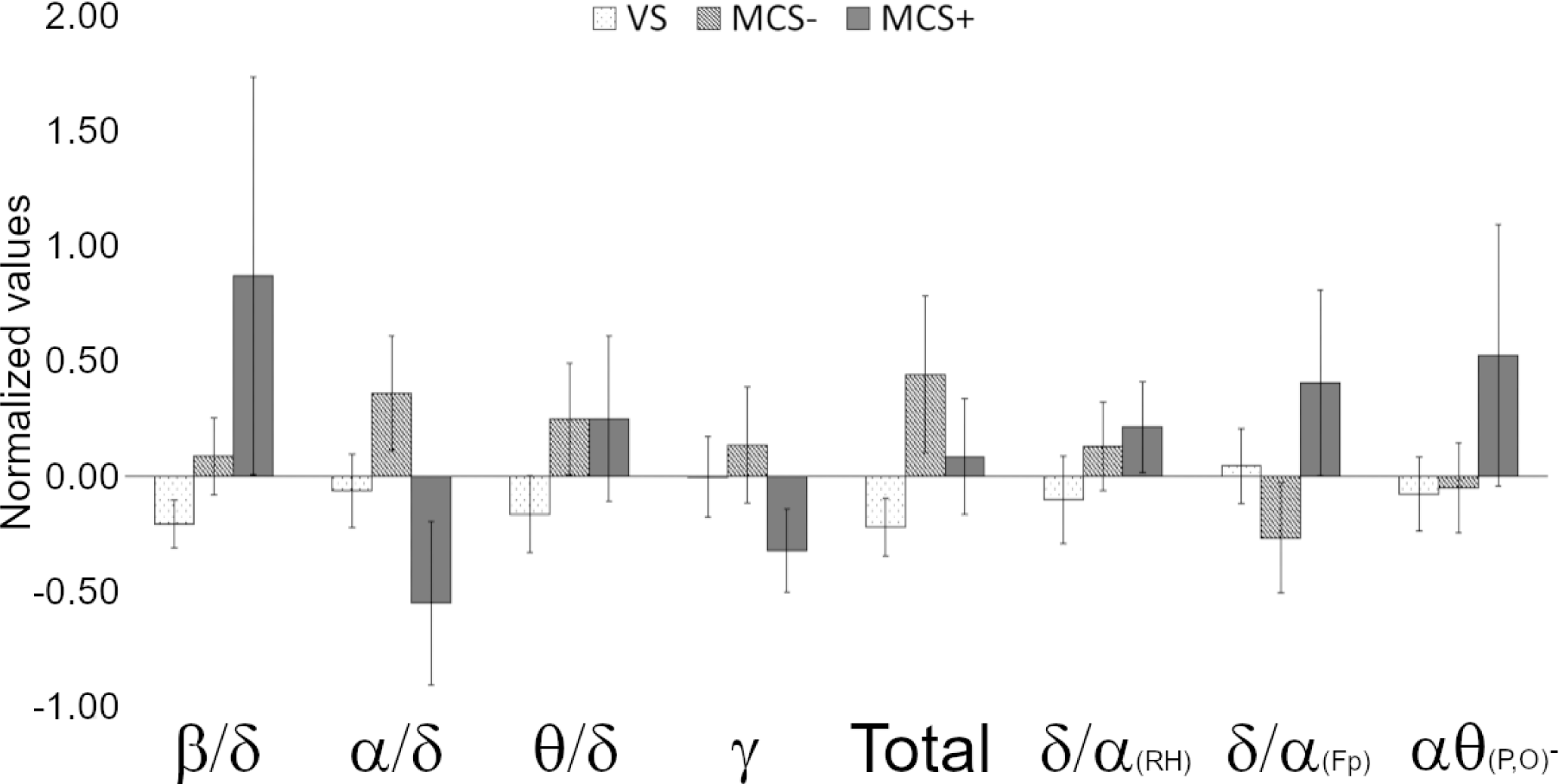
Mean intensity of each EEG component across the three DOC clinical categories. (Error bars show standard error.).

